# Forecasting next-year change in antimicrobial resistance genes in *Klebsiella pneumoniae* and *Acinetobacter baumannii* from genomic surveillance data

**DOI:** 10.64898/2026.10.10.758085

**Authors:** Tolulope S. Adeyina, Damani Andre, Ali Aslam, Kenneth Onyedibe, Muhammed Idris

## Abstract

**Background:** Surveillance tells laboratories which antimicrobial resistance genes are circulating now, but not which will change next. We built a model that forecasts, for each resistance gene and year, whether the following year brings a previously unseen allele (sequence variant), called emergence, or a change in the most common allele, called turnover. We tested it separately in *Klebsiella pneumoniae* and *Acinetobacter baumannii*.

**Results:** We used genomes annotated with AMRFinderPlus: 2,385 gene-years across 331 genes for *K. pneumoniae* and 1,059 across 216 for *A. baumannii*. To remove the effect of sequencing effort, each gene-year was sampled to 10 records and the analysis repeated 25 times. Models were trained on earlier years and tested on later ones, and on genes held out of training. On later years every model ranked genes by risk better than simple baselines in both organisms, and results for held-out genes were close. In *K. pneumoniae*, turnover was predicted better than emergence (best area under the ROC curve, AUROC, 0.909 versus 0.819). In *A. baumannii* the order reversed (emergence 0.789, turnover 0.722). The two events also relied on different features. Emergence depended mainly on how many alleles a gene had already accumulated, in both organisms. Turnover in *K. pneumoniae* depended mainly on how dominant the leading allele already was, and a stability index added a small gain. In *A. baumannii* it was spread across many weaker features. There, 56% of changes in the leading allele involved only alleles already seen, against 23% in *K. pneumoniae*, which may make them harder to predict.

**Conclusions:** Changes in resistance genes can be forecast a year ahead from routine genomic surveillance data, including for genes the model has not seen. How well it works, and what it depends on, differs between organisms, so models should be built and validated for each organism, not assumed to transfer. Such models could help laboratories decide which resistance genes to monitor most closely.

## Background

Bacterial antimicrobial resistance (AMR) was directly responsible for an estimated 1.27 million deaths in 2019 and was associated with 4.95 million [1]. *Klebsiella pneumoniae* and *Acinetobacter baumannii* were both among the six leading pathogens, which together accounted for 929,000 of the 1.27 million attributable deaths [1]. The World Health Organization lists carbapenem-resistant *A. baumannii*, and carbapenem-resistant and third-generation cephalosporin-resistant Enterobacterales (the order that includes *K. pneumoniae*), in its critical priority group [2]. *K. pneumoniae* carries a broad range of β-lactamase, aminoglycoside-modifying and efflux determinants, often including carbapenem resistance [3]. Simulations suggest that unchecked AMR could cause substantial economic damage through 2050, with annual costs comparable to those of the 2008 global financial crisis [4].

Genomic surveillance has changed how this burden is described. Public repositories hold hundreds of thousands of bacterial assemblies, and tools such as AMRFinderPlus and the Resistance Gene Identifier of CARD annotate resistance determinants at scale [5,6]. NCBI’s Pathogen Detection system and its MicroBIGG-E browser make these annotations publicly searchable across large, multi-year isolate collections [7].

Machine learning on these data has focused almost exclusively on one task: predicting an isolate’s current resistance phenotype from its genome. Models that classify isolates as resistant or susceptible, or predict minimum inhibitory concentrations, have been reported for *K. pneumoniae* and most other priority pathogens, using gene presence or absence, k-mer profiles or variant-level features [8–10]. These are diagnostic models. They answer “is this isolate resistant?”, a question about the present.

Surveillance, however, looks forward. The useful question is which resistance determinants are diversifying, and where the next novel allele is likely to appear. That question has received far less attention. Work on resistance evolution is mostly experimental, such as deep mutational scanning of single genes to map their fitness landscapes [11]. These approaches give deep insight into one gene at a time, but they do not scale to the hundreds of determinants circulating in a species, and they depend on laboratory measurement rather than on the surveillance data already being generated. The closest analogues are in virology, for example models that predict the evolution of influenza from one year to the next [12].

Forecasting first requires a different unit of analysis. An isolate is observed once and has no future, so we use the gene-year instead: a resistance determinant, in one species, in one calendar year, described by the set of sequences observed for it. At that unit, “what happens next year” has a clear meaning.

This raises a statistical problem. A gene-year is a set of alleles, and the chance of seeing a previously unseen allele rises steeply with the number of isolates sequenced (see Methods, Control of sequencing effort). A model trained on unadjusted counts can look accurate simply by detecting sequencing effort. Related problems, including population structure, clonal expansion and biased sampling, have been shown to confound machine-learning prediction of resistance across five WHO priority pathogens, and phylogeny-aware baselines and grouped validation have been recommended alongside any model [13].

Here we develop and evaluate a gene-year forecasting framework for AMR determinants in *K. pneumoniae* and *A. baumannii*, analysing each organism separately. We define two outcomes, the emergence of a previously unobserved allele and the turnover of the dominant allele, and model them separately because they describe different events. Sequencing effort is controlled by rarefying every gene-year to a fixed depth and repeating the analysis across 25 subsampling replicates (Figure 1). We ask four questions. Can each outcome be forecast a year ahead from routine surveillance data? Does the forecast beat a naive rule based on a gene’s own history? Does it hold for resistance genes absent from training?

**Figure 1.**
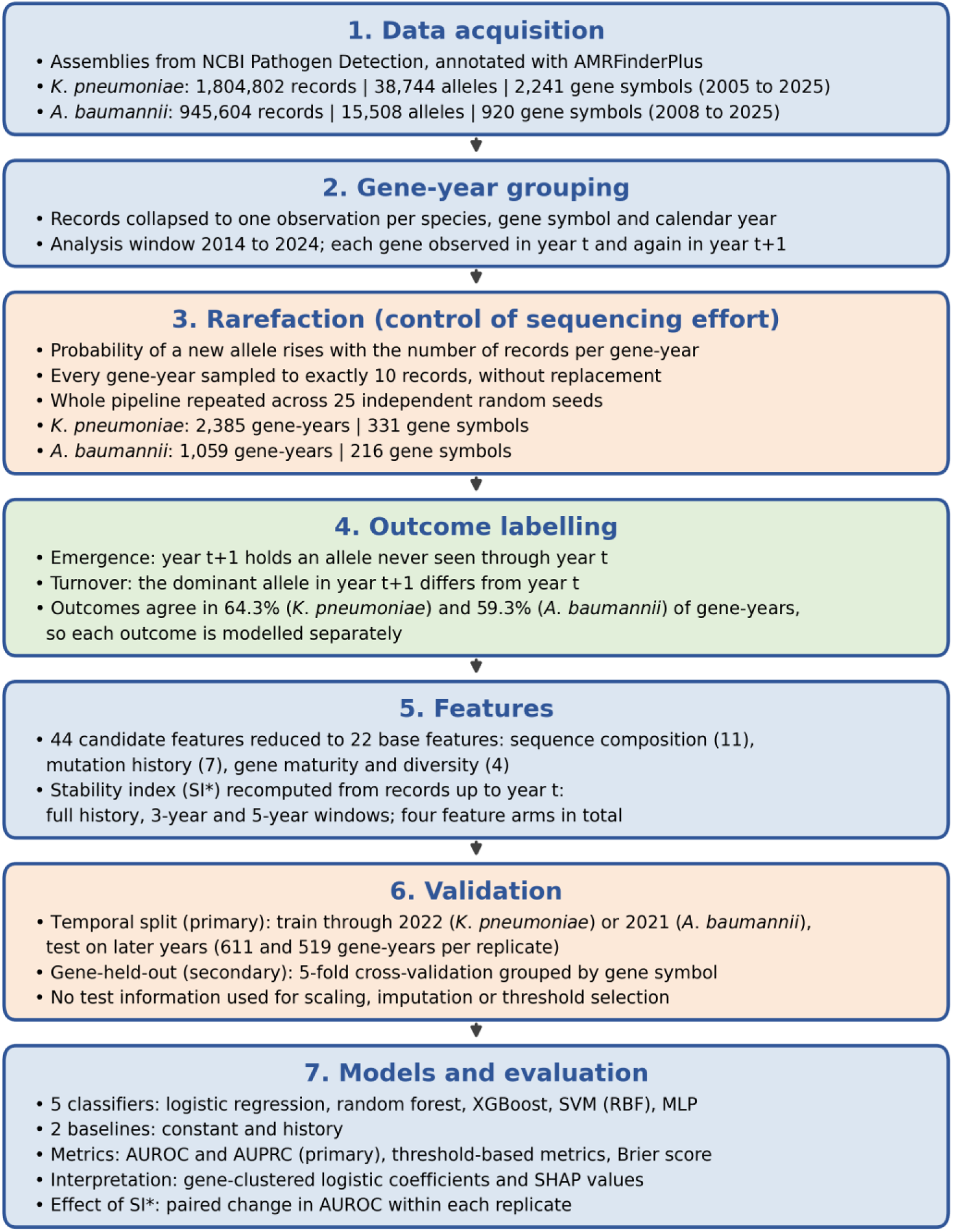
Study workflow. Resistance records annotated with AMRFinderPlus are grouped into gene-years, sampled to 10 records per gene-year across 25 replicates, labelled with two outcomes, described by 22 base features plus time-restricted stability index variants, and evaluated under two validation schemes. Counts are given for *K. pneumoniae* and *A. baumannii*. SI*, stability index; AUROC, area under the receiver operating characteristic curve; AUPRC, area under the precision-recall curve; SHAP, SHapley Additive exPlanations; MLP, multilayer perceptron; SVM, support vector machine.

And does a gene-level stability index, restricted to information available at the time of prediction, add predictive information? Running the framework in two organisms also shows which answers are specific to one species.

## Results

### The two organisms differ in panel size and in which event is more common

We began by describing the material each organism provides and how often each event occurs, since both determine what a good forecast looks like. For each organism we built a gene-year panel from AMRFinderPlus-annotated records, sampled to 10 records per gene-year (see Methods), and counted gene-years and events by year in replicate 0 (Table 1).

**Table 1.** Gene-years and event frequencies by year. Number of gene-years in each panel and the number (percentage) followed by emergence of a previously unseen allele or by turnover of the dominant allele in the next year, in replicate 0. KP, *K. pneumoniae*; AB, *A. baumannii*.

| Year | KP gene-years | KP emergence, n (%) | KP turnover, n (%) | AB gene-years | AB emergence, n (%) | AB turnover, n (%) |
| --- | --- | --- | --- | --- | --- | --- |
| 2014 | 82 | 53 (64.6) | 17 (20.7) | 40 | 17 (42.5) | 24 (60.0) |
| 2015 | 127 | 84 (66.1) | 32 (25.2) | 45 | 18 (40.0) | 28 (62.2) |
| 2016 | 165 | 92 (55.8) | 31 (18.8) | 55 | 20 (36.4) | 24 (43.6) |
| 2017 | 207 | 103 (49.8) | 41 (19.8) | 60 | 27 (45.0) | 43 (71.7) |
| 2018 | 214 | 97 (45.3) | 42 (19.6) | 63 | 24 (38.1) | 36 (57.1) |
| 2019 | 212 | 108 (50.9) | 43 (20.3) | 67 | 21 (31.3) | 34 (50.7) |
| 2020 | 233 | 108 (46.4) | 35 (15.0) | 83 | 30 (36.1) | 55 (66.3) |
| 2021 | 260 | 99 (38.1) | 41 (15.8) | 127 | 46 (36.2) | 47 (37.0) |
| 2022 | 274 | 108 (39.4) | 47 (17.2) | 150 | 65 (43.3) | 56 (37.3) |
| 2023 | 295 | 122 (41.4) | 51 (17.3) | 174 | 53 (30.5) | 65 (37.4) |
| 2024 | 316 | 107 (33.9) | 47 (14.9) | 195 | 53 (27.2) | 63 (32.3) |
| Total | 2385 | 1081 (45.3) | 427 (17.9) | 1059 | 374 (35.3) | 475 (44.9) |

The *K. pneumoniae* panel holds 2,385 gene-years across 331 genes (2014 to 2024) and the *A. baumannii* panel 1,059 gene-years across 216 genes. Both grew over time, from 82 to 316 gene-years per year in *K. pneumoniae* and from 40 to 195 in *A. baumannii*. A previously unseen allele appeared the following year in 45.3% of *K. pneumoniae* gene-years and 35.3% of *A. baumannii* gene-years, whereas the most common allele changed in 17.9% and 44.9%. Emergence was therefore the more common event in *K. pneumoniae*, and turnover in *A. baumannii*.

Because the organisms differ in size and in event frequency, a raw accuracy score cannot be compared directly between them. We therefore judge each forecast against the rate expected by chance for that event and organism.

### Emergence and turnover are distinct events

If emergence and turnover measured the same thing, they would almost always occur together and a single model would be enough. We tested this by cross-tabulating the two events within each panel (replicate 0).

The events agreed in 64.3% of *K. pneumoniae* gene-years and 59.3% of *A. baumannii* gene-years. In *K. pneumoniae*, 753 gene-years (31.6%) gained a new allele without a change in the leading allele, and 99 (4.2%) changed leader without a new allele. In *A. baumannii* the corresponding figures were 165 (15.6%) and 266 (25.1%).

Neither event is a stand-in for the other, so we modelled them separately. The organisms already differ here: changing leader without a new allele is rare in *K. pneumoniae* and common in *A. baumannii*.

### Both events can be forecast a year ahead, better than simple rules

We asked whether a gene’s own history and sequence hold enough information to forecast next year’s events better than simple rules. We trained five classifiers on gene-years up to a split year (2022 for *K. pneumoniae*, 2021 for *A. baumannii*) and tested them on later gene-years (611 and 519 per replicate), repeating the procedure over 25 sampling replicates. Each classifier was compared with two baselines: the event rate in the training years, and the gene’s own recent mutation rate (the history baseline). We report the area under the ROC curve (AUROC, where 0.5 is chance) and the area under the precision-recall curve (AUPRC), the latter judged against the event rate in the test years (Table 2, Figure 2).

**Figure 2.**
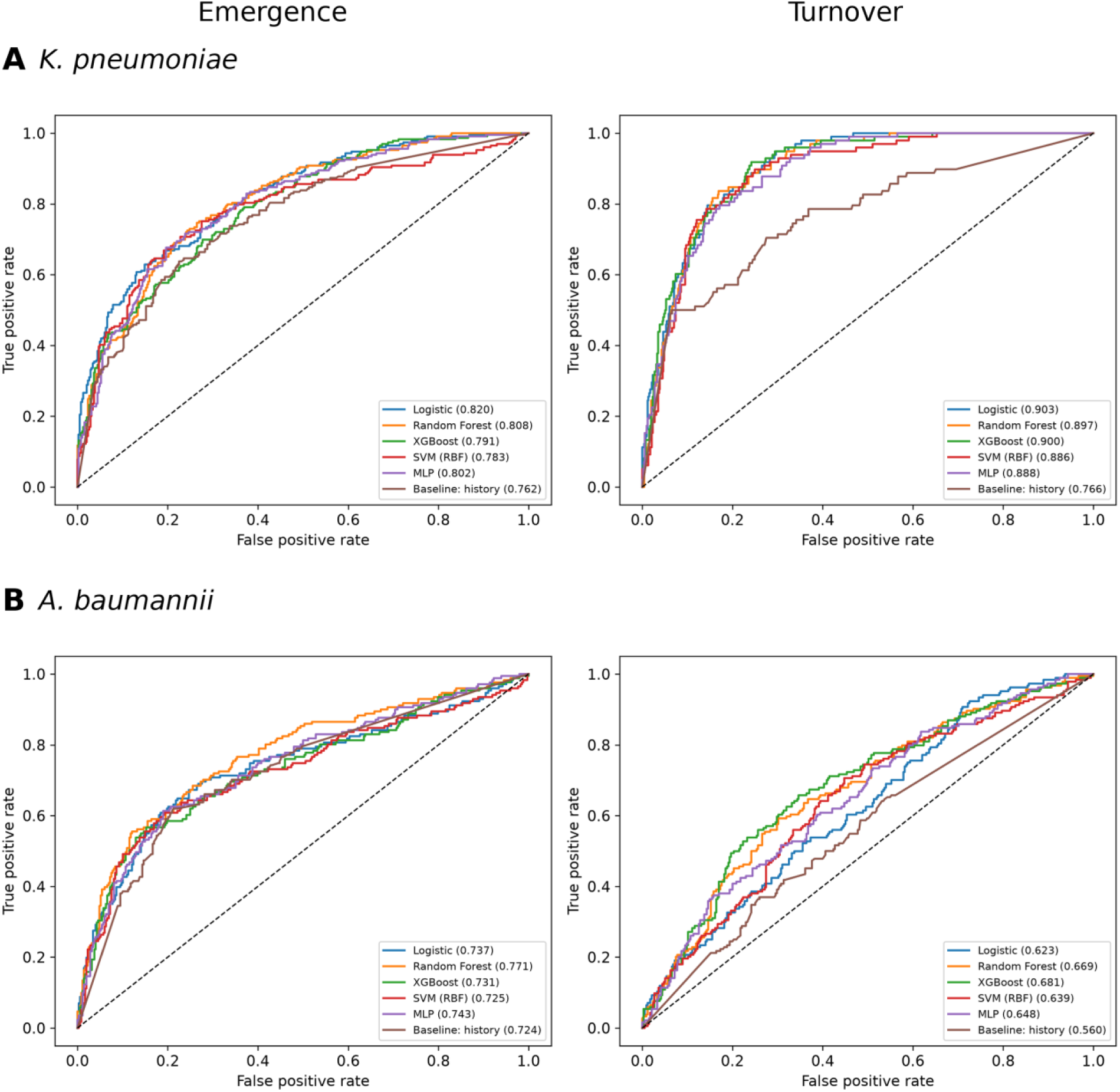
Receiver operating characteristic curves on later years. Curves for emergence (left) and turnover (right) from the temporal split in replicate 0, for the five classifiers with the full-history stability index and for the history baseline. A, *K. pneumoniae*; B, *A. baumannii*. Values in the keys are AUROC for replicate 0, and the dashed diagonal marks chance. Replicate 0 is a single draw, so its values differ from the 25-replicate means in Table 2, by more for *A. baumannii* turnover, where replicate 0 is a weaker draw than average.

**Table 2.**
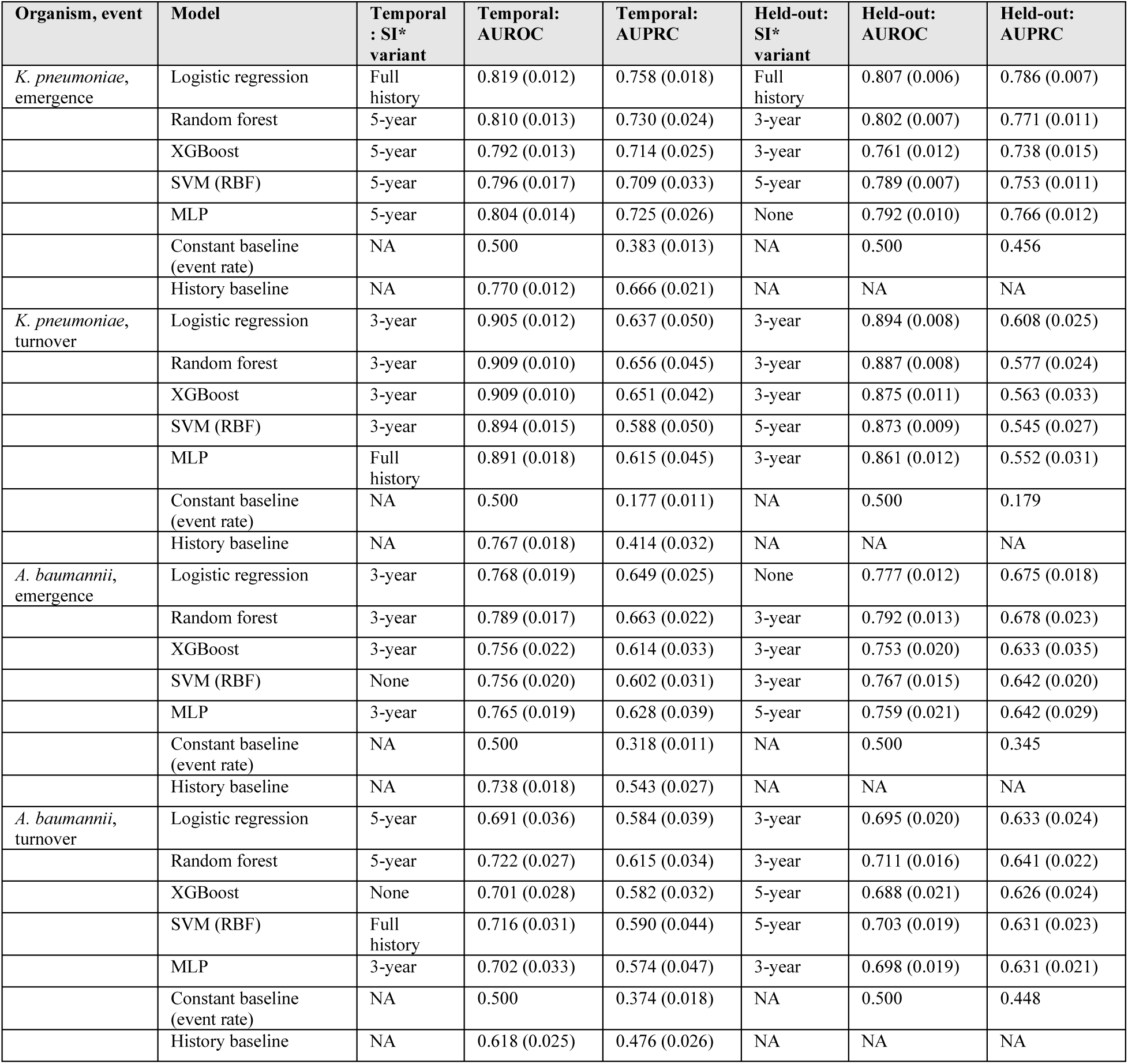
Forecasting performance on later years and on held-out genes. Mean (SD) across 25 replicates. For each model, the stability index variant shown is the one with the highest mean AUROC in that evaluation scheme. The constant baseline predicts the event rate, which gives an AUROC of 0.5 and an AUPRC equal to the event rate in the test data. The history baseline is the gene’s own recent mutation rate and was computed for later years only. NA, not applicable.

In *K. pneumoniae*, all models outperformed both baselines. Logistic regression achieved the best emergence prediction performance (AUROC 0.819 ± 0.012; AUPRC 0.758), whereas a random forest performed best for turnover prediction (AUROC 0.909 ± 0.010; AUPRC 0.656). In *A. baumannii*, random forests achieved the highest performance for both emergence (AUROC 0.789 ± 0.017; AUPRC 0.663) and turnover prediction (AUROC 0.722 ± 0.027; AUPRC 0.615). Full comparisons with chance and history-based baselines are shown in Table 2.

A gene’s history and sequence therefore carry information about next year’s events in both organisms. Emergence was predicted about equally well in the two (2.0 and 2.1 times chance), but turnover was predicted far better in *K. pneumoniae* (3.7 times chance) than in *A. baumannii* (1.6 times). The difference between the organisms in which event is easier to forecast therefore comes from turnover.

### Forecasts hold for genes withheld from training

A model that only memorised each gene’s past would fail on a gene it had never seen, so we asked whether performance holds for genes withheld from training. We repeated the evaluation with five-fold cross-validation in which all records of a gene fall in the same fold, pooling all years. This scheme has no history baseline, so models are judged against the event rate (Table 2).

Performance on held-out genes was close to that on later years. In *K. pneumoniae* the best AUROC was 0.807 for emergence and 0.894 for turnover (AUPRC 1.7 and 3.4 times chance), and in *A. baumannii* 0.792 and 0.711 (2.0 and 1.4 times chance).

The models therefore learn patterns that carry across genes, not only each gene’s own record.

### Emergence and turnover depend on different features, and turnover differs between organisms

If emergence and turnover are different events, they should depend on different features, and the organisms may differ. We examined what the forecasts relied on in two ways. First, a standardised logistic regression fitted to the full panel of each replicate, with errors clustered by gene, gave an odds ratio per standard deviation for each feature (Table 3, Figure S1). Second, SHAP values from a gradient-boosted model, fitted to the training years of replicate 0, gave each feature’s average contribution to predictions on the test years (Figure 3, Figure S2).

**Figure 3.**
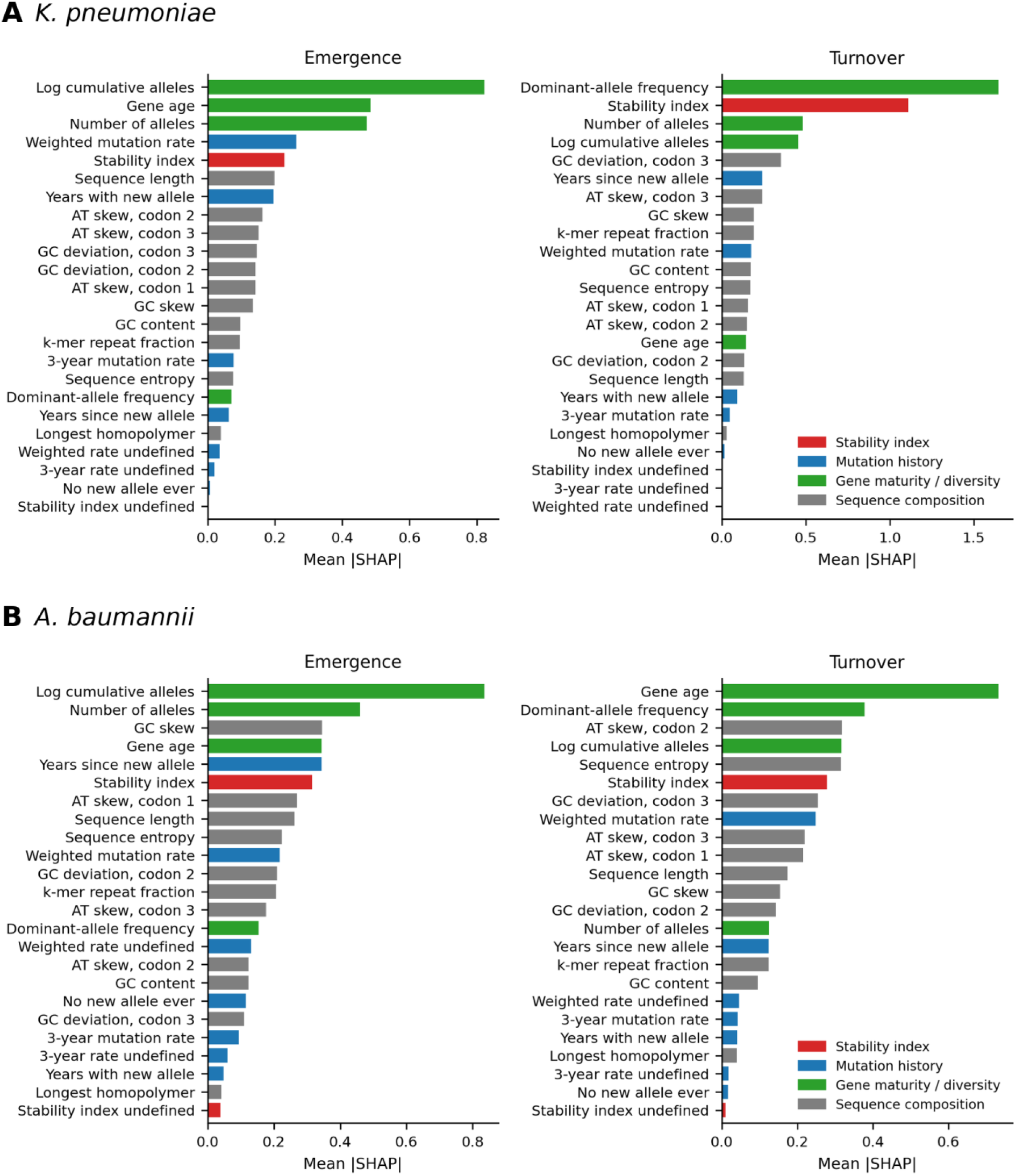
What the forecasts depend on. Mean absolute SHAP value of each feature for emergence (left) and turnover (right), from an XGBoost model fitted to the training years of replicate 0 and evaluated on the test years. A, *K. pneumoniae*; B, *A. baumannii*. Bars are coloured by feature block (stability index, mutation history, gene maturity and diversity, sequence composition), and features are sorted by importance within each panel.

**Table 3.** Logistic regression odds ratios. Odds ratio per standard deviation (mean p across 25 replicates) from a standardised logistic regression with errors clustered by gene, fitted to the full panel with the 22 base features and the full-history stability index and its missingness indicator. Features are ordered by the overall strength of their association in *K. pneumoniae*. Estimates for missingness indicators are unstable when the indicator is rare (for example 0.00 and 1,170 for *K. pneumoniae* turnover) and should not be interpreted. KP, *K. pneumoniae*; AB, *A. baumannii*.

| Feature | KP emergence | KP turnover | AB emergence | AB turnover |
| --- | --- | --- | --- | --- |
| Log cumulative allele count | 6.08 (<0.001) | 1.86 (0.089) | 3.27 (<0.001) | 1.46 (0.218) |
| Dominant-allele frequency | 0.73 (0.090) | 0.20 (<0.001) | 1.12 (0.452) | 0.70 (0.012) |
| Stability index (full history) | 1.39 (0.032) | 0.18 (<0.001) | 1.05 (0.484) | 0.76 (0.065) |
| Years with a new allele | 0.61 (0.046) | 0.50 (0.122) | 0.68 (0.187) | 0.80 (0.428) |
| 3-year mutation rate undefined (indicator) | 2.61 (0.049) | 0.00 (0.538) | 1.15 (0.472) | 1.10 (0.519) |
| Gene age | 0.65 (0.048) | 1.60 (0.268) | 0.97 (0.522) | 0.78 (0.339) |
| GC deviation, codon position 2 | 0.81 (0.109) | 1.24 (0.241) | 0.84 (0.274) | 0.98 (0.532) |
| Number of alleles | 0.93 (0.452) | 0.62 (0.031) | 1.44 (0.145) | 1.23 (0.350) |
| GC content | 0.86 (0.283) | 0.72 (0.210) | 0.75 (0.141) | 0.87 (0.454) |
| AT skew, codon position 2 | 0.91 (0.168) | 1.08 (0.476) | 1.00 (0.496) | 1.12 (0.285) |
| Sequence length | 1.08 (0.254) | 1.07 (0.323) | 1.11 (0.311) | 0.90 (0.335) |
| Years since last new allele | 0.94 (0.509) | 0.60 (0.182) | 0.78 (0.346) | 0.86 (0.433) |
| GC deviation, codon position 3 | 0.96 (0.598) | 1.37 (0.126) | 1.12 (0.377) | 1.35 (0.073) |
| Weighted mutation rate undefined (indicator) | 1.15 (0.169) | 1.07 (0.509) | 1.38 (0.047) | 0.97 (0.637) |
| AT skew, codon position 3 | 1.11 (0.122) | 1.02 (0.553) | 1.15 (0.279) | 1.00 (0.583) |
| Stability index undefined (indicator) | 0.57 (0.269) | 1,170 (0.595) | 0.81 (0.447) | 0.66 (0.270) |
| Sequence entropy | 0.95 (0.417) | 1.06 (0.485) | 0.94 (0.515) | 0.92 (0.420) |
| k-mer repeat fraction | 0.95 (0.541) | 0.86 (0.439) | 0.70 (0.160) | 0.83 (0.375) |
| No new allele ever seen | 1.01 (0.475) | 1.34 (0.418) | 1.19 (0.477) | 0.90 (0.536) |
| GC skew | 1.05 (0.474) | 0.97 (0.458) | 1.08 (0.513) | 0.99 (0.611) |
| 3-year mutation rate | 1.07 (0.486) | 0.89 (0.512) | 0.99 (0.685) | 0.87 (0.358) |
| AT skew, codon position 1 | 0.96 (0.479) | 0.98 (0.489) | 0.96 (0.594) | 0.86 (0.195) |
| Weighted mutation rate | 1.03 (0.575) | 0.98 (0.478) | 1.29 (0.222) | 0.95 (0.538) |
| Longest homopolymer | 0.99 (0.508) | 0.98 (0.626) | 1.07 (0.488) | 1.11 (0.385) |

For emergence, cumulative allele count was the strongest feature in both organisms (odds ratio 6.08 in *K. pneumoniae* and 3.27 in *A. baumannii*, and the largest SHAP value in both). Genes that had already accumulated many alleles were more likely to gain another.

For turnover in *K. pneumoniae*, dominant-allele frequency (odds ratio 0.20) and the stability index (0.18) were the strongest features: the more entrenched the leading allele, the less likely it was to be replaced. Together with allele count, the top three features carried 51% of total SHAP attribution. In *A. baumannii*, turnover was spread over many features. Gene age had the largest SHAP value (0.73), ahead of dominant-allele frequency (0.38), and the top three features carried 32% of attribution. Dominant-allele frequency was still protective (odds ratio 0.70), but the stability index was only borderline (0.76, mean p = 0.065) and ranked sixth. Sequence-composition features carried 46% of the attribution for turnover in *A. baumannii*, against 30% in *K. pneumoniae*.

The two events do depend on different things. Emergence follows how large a gene’s allele catalogue already is, in both organisms. Turnover follows how entrenched the leading allele is in *K. pneumoniae*, but has no single dominant feature in *A. baumannii*. These attributions describe what the forecasts rely on, not what causes the changes, and the SHAP values come from a single replicate.

### The stability index adds a modest gain for turnover in *K. pneumoniae* only

The stability index summarises how strongly one allele dominates a gene. Because turnover is the loss of that dominance, we expected the index to help forecast turnover but not emergence. We computed three versions from records available at each prediction year: all years so far, and trailing windows of three and five years. We added each version to each model and measured the change in AUROC within every replicate, under both validation schemes (Table 4, Figure 4). We report the mean change and the range between the 2.5th and 97.5th percentiles across replicates, which shows sensitivity to sampling and is not a confidence interval. We also compared curves with DeLong’s test in replicate 0.

**Figure 4.**
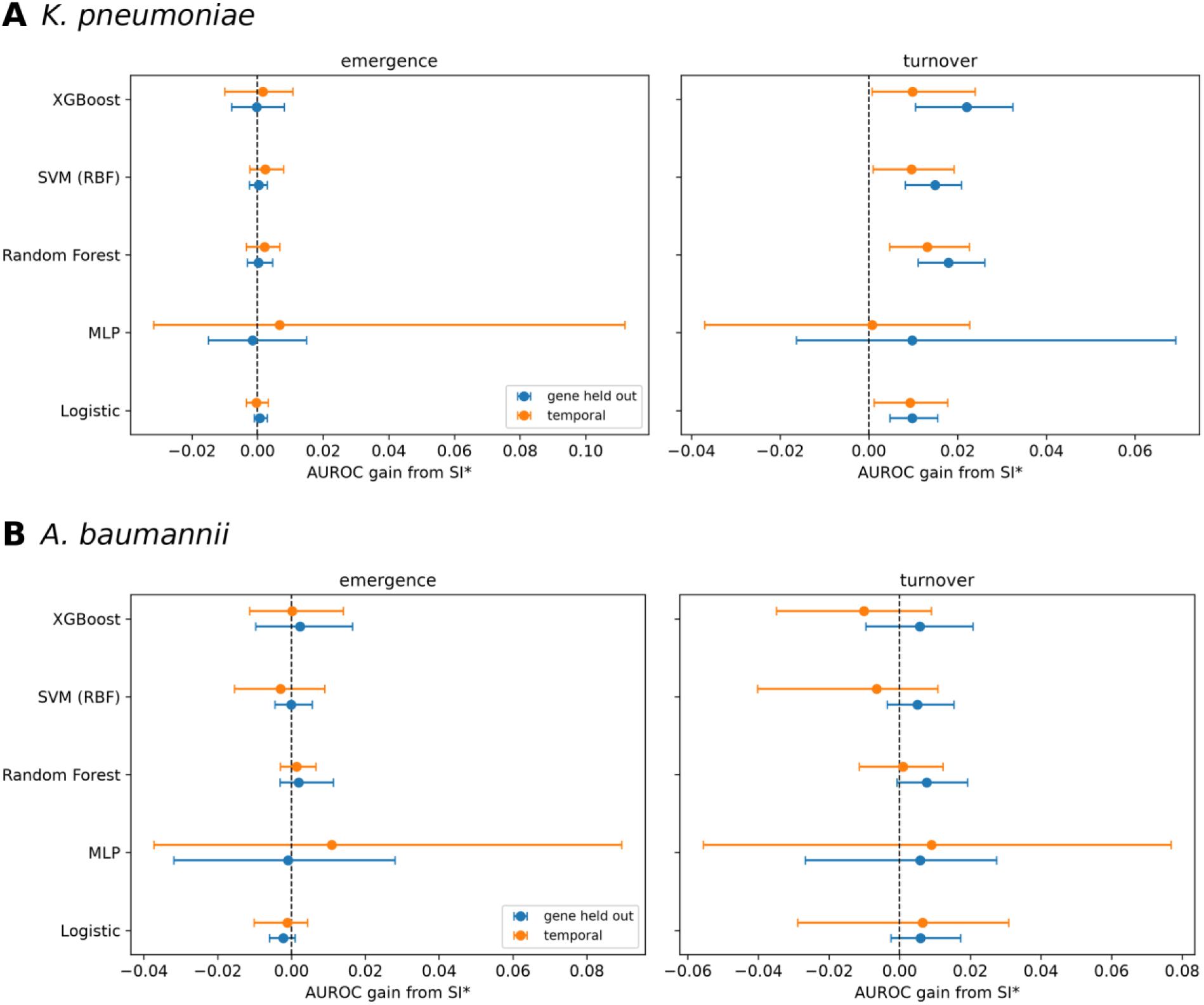
Effect of adding the stability index on AUROC. Change in AUROC when the stability index is added, averaged over its three variants, for each classifier and for emergence (left) and turnover (right), on later years (orange) and on held-out genes (blue). A, *K. pneumoniae*; B, *A. baumannii*. Points are means across 25 replicates and bars span the 2.5th to 97.5th percentile across replicates, which shows sensitivity to sampling and is not a confidence interval. The dashed line marks no change. Results for each variant are in Table 4.

**Table 4.** Change in AUROC when the stability index is added. Mean change (2.5th and 97.5th percentiles across 25 replicates) for each classifier and stability index variant, on later years and on held-out genes. Positive values mean the index improved AUROC. The ranges describe sensitivity to sampling and are not confidence intervals.

| Organism, event | Model | Temporal: 3-year | Temporal: 5-year | Temporal: full history | Held-out: 3-year | Held-out: 5-year | Held-out: full history |
| --- | --- | --- | --- | --- | --- | --- | --- |
| *K. pneumoniae*, turnover | Logistic regression | +0.010 (+0.002, +0.018) | +0.010 (+0.003, +0.018) | +0.008 (0.000, +0.016) | +0.010 (+0.005, +0.015) | +0.011 (+0.005, +0.016) | +0.009 (+0.004, +0.015) |
|  | Random forest | +0.015 (+0.009, +0.024) | +0.013 (+0.004, +0.021) | +0.011 (+0.005, +0.018) | +0.020 (+0.012, +0.029) | +0.018 (+0.012, +0.025) | +0.016 (+0.011, +0.025) |
|  | XGBoost | +0.011 (+0.001, +0.024) | +0.010 (+0.001, +0.024) | +0.008 (+0.003, +0.018) | +0.023 (+0.011, +0.032) | +0.023 (+0.012, +0.033) | +0.020 (+0.010, +0.030) |
|  | SVM (RBF) | +0.011 (+0.001, +0.021) | +0.010 (+0.001, +0.017) | +0.007 (+0.002, +0.014) | +0.015 (+0.008, +0.020) | +0.016 (+0.010, +0.021) | +0.014 (+0.009, +0.019) |
|  | MLP | -0.002 (-0.041, +0.023) | +0.001 (-0.026, +0.023) | +0.003 (-0.023, +0.019) | +0.010 (-0.015, +0.060) | +0.010 (-0.014, +0.051) | +0.010 (-0.016, +0.049) |
| *K. pneumoniae*, emergence | Logistic regression | -0.001 (-0.004, +0.002) | -0.001 (-0.003, +0.001) | +0.001 (-0.003, +0.004) | +0.000 (-0.001, +0.002) | +0.001 (-0.001, +0.002) | +0.001 (0.000, +0.004) |
|  | Random forest | +0.002 (-0.003, +0.006) | +0.003 (-0.004, +0.008) | +0.002 (-0.003, +0.005) | +0.000 (-0.004, +0.004) | +0.001 (-0.003, +0.004) | 0.000 (-0.003, +0.004) |
|  | XGBoost | +0.001 (-0.010, +0.010) | +0.002 (-0.010, +0.012) | +0.002 (-0.004, +0.010) | +0.001 (-0.003, +0.008) | 0.000 (-0.008, +0.006) | -0.002 (-0.008, +0.007) |
|  | SVM (RBF) | +0.002 (-0.003, +0.007) | +0.003 (-0.001, +0.010) | +0.002 (-0.001, +0.007) | +0.000 (-0.002, +0.003) | +0.001 (-0.002, +0.003) | 0.000 (-0.003, +0.002) |
|  | MLP | +0.005 (-0.035, +0.112) | +0.008 (-0.020, +0.111) | +0.007 (-0.021, +0.110) | -0.002 (-0.017, +0.013) | -0.001 (-0.015, +0.015) | -0.002 (-0.012, +0.013) |
| *A. baumannii*, turnover | Logistic regression | +0.001 (-0.032, +0.029) | +0.011 (-0.005, +0.028) | +0.007 (-0.010, +0.028) | +0.006 (-0.002, +0.021) | +0.006 (-0.003, +0.020) | +0.005 (-0.002, +0.015) |
|  | Random forest | -0.003 (-0.015, +0.006) | +0.003 (-0.008, +0.013) | +0.003 (-0.008, +0.012) | +0.009 (+0.001, +0.020) | +0.008 (+0.000, +0.017) | +0.006 (-0.003, +0.016) |
|  | XGBoost | -0.019 (-0.042, +0.000) | -0.006 (-0.022, +0.010) | -0.005 (-0.021, +0.010) | +0.005 (-0.009, +0.015) | +0.008 (-0.008, +0.025) | +0.004 (-0.009, +0.016) |
|  | SVM (RBF) | -0.019 (-0.046, +0.004) | -0.002 (-0.013, +0.011) | +0.001 (-0.013, +0.013) | +0.005 (-0.004, +0.016) | +0.006 (-0.003, +0.016) | +0.004 (-0.001, +0.011) |
|  | MLP | +0.011 (-0.034, +0.070) | +0.012 (-0.052, +0.069) | +0.004 (-0.072, +0.066) | +0.005 (-0.021, +0.025) | +0.006 (-0.024, +0.027) | +0.006 (-0.033, +0.025) |
| *A. baumannii*, emergence | Logistic regression | -0.001 (-0.008, +0.005) | -0.002 (-0.010, +0.002) | -0.001 (-0.005, +0.003) | -0.002 (-0.005, +0.001) | -0.004 (-0.007, -0.001) | -0.002 (-0.004, +0.001) |
|  | Random forest | +0.002 (-0.002, +0.005) | +0.001 (-0.003, +0.006) | +0.001 (-0.003, +0.007) | +0.002 (-0.001, +0.008) | +0.002 (-0.003, +0.008) | +0.002 (-0.002, +0.008) |
|  | XGBoost | +0.002 (-0.010, +0.015) | +0.001 (-0.011, +0.012) | -0.002 (-0.017, +0.010) | +0.005 (-0.009, +0.015) | -0.001 (-0.010, +0.013) | +0.002 (-0.009, +0.015) |
|  | SVM (RBF) | -0.002 (-0.013, +0.012) | -0.004 (-0.017, +0.004) | -0.003 (-0.015, +0.006) | +0.001 (-0.004, +0.006) | -0.001 (-0.004, +0.005) | 0.000 (-0.005, +0.006) |
|  | MLP | +0.011 (-0.029, +0.071) | +0.012 (-0.030, +0.075) | +0.010 (-0.034, +0.070) | -0.001 (-0.027, +0.030) | +0.001 (-0.027, +0.026) | -0.003 (-0.037, +0.020) |

In *K. pneumoniae*, adding the index changed turnover AUROC by −0.002 to +0.015 on later years (an increase in 14 of 15 combinations of model and version) and by +0.009 to +0.023 on held-out genes (15 of 15). The replicate range excluded zero in 11 of 15 and 12 of 15 combinations. The exceptions were the neural network in every combination, and logistic regression with the all-years version on later years. For emergence the changes were within 0.01 and none of the 30 ranges excluded zero. The three versions gave similar results. DeLong tests on later years were significant in 8 of 15 turnover comparisons (random forest and XGBoost with all three versions, and the SVM with two) and in 4 of 15 emergence comparisons.

In *A. baumannii*, turnover AUROC changed by −0.019 to +0.012 on later years (an increase in 9 of 15 combinations, with no range excluding zero) and by +0.004 to +0.009 on held-out genes (an increase in all 15, but only 2 of 15 ranges excluded zero). For emergence the changes ranged from −0.004 to +0.012 and from −0.004 to +0.005. One range lay wholly below zero (logistic regression with the 5-year version, on held-out genes) and none lay above it. DeLong tests were significant in 6 of 15 turnover and 5 of 15 emergence comparisons, but in most of them (4 of 6 and 3 of 5) the index lowered AUROC.

The expectation held for *K. pneumoniae*: the index gave a modest and mostly reliable gain for turnover and none for emergence. It did not hold for *A. baumannii*, where the index added nothing consistent for either event. For emergence in *K. pneumoniae*, the direction of the association depended on the model: it was positive in the logistic regression (odds ratio 1.39, mean p = 0.032) but negative in the SHAP analysis.

### Turnover in *A. baumannii* often involves alleles already seen

Turnover was harder to predict in *A. baumannii*. We asked whether this is because the leading allele there often changes without any new allele appearing, a change that a gene’s own history gives little warning of. In replicate 0 we split turnover events by whether a new allele also appeared that year, and compared turnover frequency between training and test years. We also compared performance on held-out genes, which pools years and so removes any shift over time.

In *A. baumannii*, 266 of 475 turnover events (56%) involved no new allele, against 99 of 427 (23%) in *K. pneumoniae*. Turnover frequency in *A. baumannii* fell from 54% of gene-years in the training years to 35% in the test years, against 19% to 16% in *K. pneumoniae*. On held-out genes, where this shift does not apply, turnover remained weaker in *A. baumannii* (AUROC 0.711 against 0.894).

These findings are consistent with the idea that much of the turnover in *A. baumannii* is a change of leader among known alleles, which is harder to predict from a gene’s history. The fall in turnover frequency over time probably adds to the gap on later years, but it cannot explain it, because the gap persists on held-out genes. We did not test what drives these changes, such as lineage-driven sampling or chance swings between near-equal alleles, and return to this in the Discussion.

## Discussion

### Principal findings

We built a gene-year model that forecasts, a year ahead, whether a resistance gene will gain a new allele (emergence) or lose its leading allele (turnover), and tested it separately in *K. pneumoniae* and *A. baumannii*. In both organisms the forecasts beat simple rules, and they held for genes withheld from training. How well they worked, and what they relied on, differed between the organisms. Emergence followed the size of a gene’s allele catalogue in both and was predicted about equally well. Turnover followed how entrenched the leading allele was in *K. pneumoniae*, where it was predicted well, and was spread across many features and harder to predict in *A. baumannii*.

### Emergence follows the size of the allele catalogue

Cumulative allele count was the strongest feature for emergence in both organisms: genes that had already accumulated many alleles were more likely to gain another. One reading is that some genes keep diversifying. Another is that a larger catalogue has simply been sampled more thoroughly over time, which raises the chance that a newly sampled allele is new to the record. Rarefaction fixes sampling effort within each year but not across a gene’s history, so these data cannot separate the two. The practical gain is modest. The best emergence models improved on a gene’s own recent mutation rate by about 0.05 AUROC in both organisms (0.819 against 0.770 in *K. pneumoniae*, and 0.789 against 0.738 in *A. baumannii*), so most of the usable signal for emergence is already in a gene’s recent history.

### Why turnover differs between the organisms

In *K. pneumoniae*, turnover was predicted far better than emergence (AUROC 0.909, against 0.767 for the history baseline), and it depended mainly on dominant-allele frequency and the stability index. Most turnover events there (77%) came with a new allele, so turnover is largely an established leader being displaced by a newcomer, and how entrenched the leader is indicates how likely that is.

In *A. baumannii*, turnover was predicted less well (AUROC 0.722, against 0.618 for the history baseline), although the gain over the baseline was still large, and its predictors were spread across many features.

We first suspected that *A. baumannii* simply produces new variants less often. It does gain new alleles less often (35% of gene-years against 45%), but the larger difference lies elsewhere: 56% of its turnover events involved no new allele, against 23% in *K. pneumoniae*. The leading allele in *A. baumannii* often changes among alleles that are already known, which gives a gene’s own history little warning.

Three explanations seem plausible, and we have tested none of them. The sequenced isolates may change from year to year, with different lineages or outbreaks dominating the record, so that a different known allele leads without any new mutation. Population structure and clonal expansion are known to confound machine learning on resistance data [13]. Alternatively, at 10 records per gene-year, two alleles of near-equal frequency can swap leadership by chance. Finally, the panel itself changed: it grew from 40 gene-years in 2014 to 195 in 2024, and turnover frequency fell from 54% in the training years to 35% in the test years. This shift probably accounts for part of the weaker temporal result, but the gap remains on held-out genes (AUROC 0.711 against 0.894), where years are pooled. Lineage information for each isolate, and reruns at other sampling depths, would help separate these explanations.

### The stability index is a modest, organism-specific contributor

The stability index illustrates a design point for forecasting studies. As originally described, it is computed once from a gene’s whole record, which would leak future information if used naively, so we recomputed it from the records available at each prediction year. With that restriction it behaved differently in the two organisms. In *K. pneumoniae* it was the second most important feature for turnover, carried a strong protective association, and added a small but mostly reliable gain of about 0.01 AUROC. In *A. baumannii* its association was weak, it ranked sixth, and the gain was not consistent. For emergence it added nothing measurable in either organism, and in *K. pneumoniae* the direction of its association depended on the model.

The pattern fits how the index is built. It combines how much one allele dominates with how evenly the others are spread, so it describes how entrenched the leader is, which matters most where turnover means a leader being displaced. This is a plausible reading, not a tested mechanism. A time-restricted stability index is best seen as an interpretable summary of dominance that helped in one organism, not as a transferable predictor.

### Using the forecasts

A laboratory could use the model to rank genes by their risk of changing in the coming year and decide which to watch most closely. It needs only the annotated data that a surveillance laboratory already holds. It works for genes the model was not trained on, provided they have a few years of surveillance records. And it gives a ranking, which suits prioritisation better than a yes-or-no call. Our results also show how it should be used: the features that matter, and the accuracy to expect, differ between organisms, so a model should be built and validated for each organism and not carried over.

### Limitations

- **What is forecast is allele change, not resistance.** Alleles are defined by exact nucleotide identity, so a change may be silent and need not alter resistance or clinical impact. Linking these forecasts to phenotype is a separate step.
- **Public data are not a random sample.** Pathogen Detection reflects which isolates laboratories choose to sequence and submit. Rarefaction removes the effect of sequencing effort within a gene-year, but not this wider sampling bias.
- **Several choices were fixed in advance.** We used 10 records per gene-year, one split year per organism, and fixed model settings without tuning, and we did not test other depths. Reported performance is therefore likely a lower bound for tuned models, and results may depend on the sampling depth.
- **The replicates are not independent.** The 25 replicates are repeated samples from one panel, so their ranges describe sensitivity to sampling and not uncertainty about a population effect. The SHAP values come from a single replicate.
- **The organisms differ in panel size.** The *A. baumannii* panel is less than half the size of the *K. pneumoniae* panel, so some differences between organisms may reflect size and not biology.
- **The horizon is one year.** We did not test longer horizons.

### Conclusions

A gene-year model built from routine genomic surveillance data can forecast, a year ahead, which resistance genes will gain a new allele and which will change their leading allele, in *K. pneumoniae* and *A. baumannii*, including for genes absent from training. Emergence followed the size of a gene’s allele catalogue and was predicted about equally well in both organisms. Turnover followed how entrenched the leading allele was and was predicted well in *K. pneumoniae*, where a time-restricted stability index added a small gain, but less well in *A. baumannii*, where it often involved alleles already known. Because performance and drivers differ between organisms, forecasting models should be built and validated for each organism, not assumed to transfer. Next steps are to test the explanations for *A. baumannii* turnover with lineage information, to link allele forecasts to resistance phenotype, and to evaluate the model prospectively.

## Methods

### Study design and data source

This was a retrospective modelling study of publicly archived bacterial genome assemblies. No isolates were collected and no laboratory work was done. Predictors were built from data available up to and including a calendar year t, and outcomes were measured in year t+1. The same pipeline was applied independently to *K. pneumoniae* and *A. baumannii*, each with its own panel, split year and results, and the two were never pooled.

Genome assemblies were retrieved from the NCBI Pathogen Detection system, and resistance determinants were annotated with AMRFinderPlus. Full-length nucleotide sequences were extracted using the start and stop coordinates reported by AMRFinderPlus, with reverse complements taken for genes on the negative strand. Each record carried 14 fields: species, isolate identifier, gene symbol, gene name, resistance class and subclass, start, stop and strand, nucleotide and amino-acid sequence, sequence length, collection year, and a gene-level stability index supplied with the source file (see Stability index).

The *K. pneumoniae* input comprised 1,804,802 records, with 38,744 distinct nucleotide sequences across 2,241 gene symbols, spanning 2005 to 2025. Class and subclass were missing in 15.50% of these records; every field required for the analysis (species, gene symbol, collection year, isolate identifier, nucleotide sequence) was complete. The *A. baumannii* input comprised 945,604 records, with 15,508 distinct sequences across 920 gene symbols, spanning 2008 to 2025, and none had missing class or subclass.

Gene name, class and subclass are properties of a gene symbol. Where records of one symbol carried inconsistent values, the most frequent value was used; no records were removed.

### Unit of analysis and allele definition

The unit of analysis is the gene-year: one observation per combination of species, gene symbol and calendar year. A gene-year is described by the set of nucleotide sequences, called alleles, observed for that determinant in that year across all isolates carrying it. Two records share an allele if and only if their nucleotide sequences are identical after case normalisation, so any substitution, insertion or deletion defines a new allele. No amino-acid or similarity-threshold filtering was applied.

We aggregate to the gene-year because lag features and outcomes computed row by row over an isolate-sorted table would compare records from the same year, not from adjacent years.

### Control of sequencing effort

The probability of observing a previously unseen allele in a gene-year rises with the number of records available for it. In *K. pneumoniae* it rises from 0.37 in gene-years with five or fewer records to 0.91 in those with more than 200, and in *A. baumannii* from 0.40 to 0.80. Sequencing effort therefore confounds any diversity-derived outcome, and it varied with calendar time as surveillance broadened.

To remove this dependence, every gene-year was rarefied to exactly 10 records drawn without replacement, and gene-years with fewer than 10 records were excluded. A single subsample is one arbitrary realisation, so the entire pipeline (rarefaction, gene-year collapse, feature construction and outcome definition) was repeated across 25 random seeds. Models were fitted separately within each replicate, and every reported metric is the mean and standard deviation across the 25 replicates. Replicates were never pooled into one training set.

Rarefaction fixes the number of records per gene-year at 10, not the number of isolates, because one isolate can contribute more than one record to a gene-year, for example through a paralogous copy or a split contig. Among the 10 records, the number of unique isolates ranged from 6 to 10 in *K. pneumoniae* and from 5 to 10 in *A. baumannii*. We therefore refer to a depth of 10 records throughout.

### Panel construction

A record was eligible if it had a non-missing nucleotide sequence, collection year and gene symbol. The analysis window starts in 2014 for both organisms, the first year of contiguous coverage in *K. pneumoniae*. A gene-year entered the panel if the gene was observed at depth 10 or more in year t and again in year t+1, so that both outcomes are defined; the final calendar year has no successor and contributes no rows. Because membership depends only on record counts, the panel is identical across the 25 replicates.

The *K. pneumoniae* panel comprises 2,385 gene-years across 331 gene symbols, and the *A. baumannii* panel 1,059 gene-years across 216 gene symbols, both spanning 2014 to 2024.

### Outcome definitions

Because a gene-year is a set of alleles, “mutation in the following year” needs an explicit definition. Two binary outcomes were specified a priori. Let A(g,t) be the set of alleles observed for gene g in year t, and C(g,t) the cumulative set of alleles observed through year t.

*Emergence of a novel allele.* The outcome is positive if year t+1 contains at least one allele never previously observed for that gene. A single isolate carrying a novel sequence is sufficient.

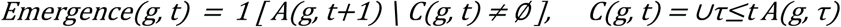

*Turnover of the dominant allele.* Let d(g,t) be the most frequent allele in year t, with ties broken by the lowest allele identifier. The outcome is positive if the prevailing sequence is displaced in the following year.

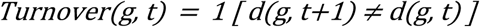

Both outcomes are defined on the same rows, namely gene-years whose successor year was observed.

### Features and feature selection

Features were computed from each gene-year’s dominant allele (sequence composition) and from the gene’s history through year t. The base set contains 22 features in three blocks.

- **Sequence composition (11).** GC content, GC skew, GC deviation at codon positions 2 and 3, AT skew at each of the three codon positions, k-mer repeat fraction, longest homopolymer, Shannon entropy of the sequence, and sequence length. Homopolymer and repeat content are substrates for slipped-strand mispairing, GC content and skew relate to replication-associated mutational asymmetry, and codon-position GC deviation helps separate mutational bias from selective constraint.
- **Mutation history (7).** The rate at which new alleles appeared over the previous three years, an exponentially weighted version of that rate (span of five years), years since the last new allele, an indicator that no new allele has ever been seen, the number of years in which a new allele appeared, and missingness indicators for the two rates.
- **Gene maturity and diversity (4).** Log cumulative allele count, gene age (years since first observation), number of alleles in the current year, and dominant-allele frequency. The size of a gene’s existing allele catalogue conditions the probability that a further novel allele is detected, a discovery-pool effect distinct from biological mutation rate, and gene age captures time under surveillance.

Feature selection used collinearity and variance only, was completed before any model was fitted, and did not use the outcomes. Of the 44 candidate features (including the two stability-index columns), we removed those that were constant, exactly determined by other members of a compositional group, near-duplicates of another feature (absolute Pearson correlation above 0.95), or restatements of GC content.

The raw codon-position nucleotide fractions were replaced by GC-deviation and AT-skew features. Calendar year was not used as a predictor, because under the temporal split its test values lie outside the training range, and gene age and cumulative allele count already capture time under surveillance.

No values were imputed. An undefined quantity, such as years since the last new allele in a gene that has never produced one, is a fact about the gene and not a gap in measurement, so each was paired with a binary indicator and a fixed fill (0 for undefined rates, and gene age plus 1 for years since the last new allele). The feature matrix contained no missing values, verified by assertion before every model fit.

### Stability index

The Stability Index (SI*) is a gene-level summary of how strongly one allele dominates, weighted by how unevenly the alleles are distributed:

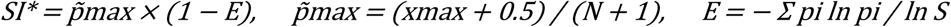

Here x_max is the count of the most frequent sequence, N the total number of sequences, S the number of distinct variants, p_i the relative frequency of variant i, and E the Shannon evenness (set to zero when only one variant has been seen). The Jeffreys-type correction stabilises the dominance estimate at small N.

The SI* column in the source file is a single value per gene computed over the complete 2005 to 2025 series, so it carries information from years after any given prediction point. We kept it for provenance only and never used it as a feature. Instead, SI* was recomputed for every gene-year from the rarefied allele counts using only records available through year t, in three forms: an expanding window over all years up to t (full history) and trailing windows of three and five years.

The index is unstable when few records contribute, so values were masked where fewer than 30 rarefied records had accumulated within the window, equivalent to at least three gene-years at depth 10. The floor was chosen from a sweep of 10, 20, 30, 40, 50, 75 and 100. In *K. pneumoniae*, a floor of 30 kept SI* defined for 76.4% of gene-years and reduced mean year-over-year volatility by 20.5% relative to no floor, with no improvement between floors of 10 and 20 and diminishing returns beyond 30. For *A. baumannii* the same floor was kept for consistency across organisms, although that organism’s own retention and volatility curve alone would have supported a higher floor of 40 to 50. Masked values were set to zero and paired with a missingness indicator, so each variant enters a model as two columns added to the 22-feature base set. This gives four feature arms: no SI*, and each of the three variants.

### Models, class imbalance and thresholds

Five classifiers were compared against two baselines. Hyperparameters were fixed a priori (Table 5) and random seeds were fixed at 0.

**Table 5.** Specification of the classifiers and baselines. Hyperparameters were fixed in advance and random seeds were set to 0.

| Model | Specification |
| --- | --- |
| Logistic regression | Standardised inputs; L2 penalty with C = 1; balanced class weights; up to 2,000 iterations |
| Random forest | 400 trees; minimum 5 samples per leaf; balanced class weights |
| XGBoost | 400 trees; maximum depth 4; learning rate 0.05; row and column subsampling 0.8; no class weighting |
| Support vector machine | RBF kernel; C = 1; gamma set to scale; standardised inputs; balanced class weights |
| Multilayer perceptron | Standardised inputs; hidden layers of 64 and 32 units; L2 penalty 0.001; early stopping with patience 20 |
| Baseline: constant | Predicts the training-partition prevalence for every gene-year |
| Baseline: history | Exponentially weighted new-allele rate (span of five years) used directly as the score |

Imbalance handling is deliberately not uniform. Class weights inversely proportional to class frequency were applied to logistic regression, random forest and the SVM. XGBoost was fitted without instance weighting, and the MLP implementation does not support class weights. No oversampling, undersampling or synthetic minority generation was used, because these would alter the outcome distribution and the prevalence-referenced reading of AUPRC. Performance is therefore summarised by AUROC and by AUPRC read against the outcome-specific no-skill floor, which equals test prevalence. No hyperparameter search was run, which keeps all models under identical conditions across 25 replicates, two outcomes and two validation schemes and prevents test information from reaching model configuration. The cost is that reported performance is likely a lower bound on what tuned models could achieve.

For each model, the classification threshold was the cutpoint that maximised F1 over out-of-fold predictions within the training partition (3-fold stratified cross-validation, 60 candidate cutpoints between the 2nd and 98th percentiles of the out-of-fold scores), applied unchanged to the test partition. If the inner cross-validation could not run, the training prevalence was used. The constant baseline classifies by the training prevalence, and the history baseline was thresholded at the median of its own test-set scores, which uses no labels. Threshold-dependent metrics for the baselines therefore follow a different rule from those of the models, so AUROC and AUPRC are the primary comparisons.

### Validation design

Two evaluation schemes were run, and they were never combined.

*Temporal split (primary).* Models were trained on gene-years up to a split year and tested on later gene-years within the same rarefaction replicate. Genes appear on both sides by design, because the task is to forecast the behaviour of determinants already under surveillance from their own accumulated history. In *K. pneumoniae* the split year was 2022, giving 611 test gene-years per replicate (2023 and 2024, with outcomes observed in 2024 and 2025) and 287 gene symbols present in both partitions. In *A. baumannii* the split year was 2021, giving 519 test gene-years per replicate (2022 to 2024) and 130 shared gene symbols. Outcomes for the final training year are measured in the first test year, but no test outcome is used in training.

The split year was chosen per organism from the row balance, year coverage and class balance on each side. For *K. pneumoniae*, an earlier candidate of 2019 put about twice as many rows in the test set as in training, and 2022 was adopted. For *A. baumannii*, 2022 gave a 65% to 35% train-to-test split with only two test years, whereas 2021 gave 51% to 49% and three test years.

*Gene-held-out cross-validation (secondary).* Five-fold grouped cross-validation with gene symbol as the grouping variable, pooling all years, so that every test gene is entirely unseen by the model during training. This evaluates generalisation to resistance genes absent from training and addresses confounding by repeated sampling of related determinants. Metrics were averaged across folds within a replicate.

Baselines were computed only for the temporal split, so under this scheme models are judged against the outcome frequency.

### Statistical analysis and interpretation

Discrimination was summarised by AUROC and AUPRC. Threshold-dependent metrics were also computed at the training-derived threshold but are not reported here.

*Effect of the stability index.* For each SI* variant, the AUROC and AUPRC difference from the no-SI* arm was computed within each replicate, for the same outcome, scheme and model, and summarised as the mean, the 2.5th to 97.5th percentile range across replicates, and the proportion of replicates with a positive difference. Replicates are repeated subsamples of one panel, so these ranges describe sensitivity to rarefaction and are not confidence intervals for a population effect. On the temporal split for replicate 0 only, DeLong’s test compared the correlated AUROC curves of each SI* arm against the no-SI* arm, and McNemar’s test compared classifications at the selected threshold (exact when discordant pairs numbered fewer than 25).

*Coefficients.* A standardised, unpenalised logistic regression was fitted to the full panel in each replicate, using the 22 base features and the full-history SI* pair, with standard errors clustered by gene symbol because each gene contributes several gene-years. Coefficients are log-odds per standard deviation; z statistics and p values are averaged across the 25 replicates.

*Feature attribution.* SHAP values were computed with TreeExplainer for an XGBoost model fitted on the temporal training partition of replicate 0 with the same feature set, and evaluated on the temporal test partition. Importance is the mean absolute SHAP value (log-odds scale) per feature, summed within feature blocks. Because this uses a single replicate, attribution is treated as qualitative.

### Prevention of information leakage

- Every feature, including the stability index (see Stability index), is a function of data at or before year t. Rolling windows end at t, and the cumulative allele set that defines emergence excludes year t+1.
- Standardisation is performed inside fitted pipelines, so scaling parameters come from training data only.
- Thresholds were selected on training data only, except for the label-free median rule for the history baseline described above.
- Non-independence of gene-years from the same gene is addressed by gene-held-out validation and by gene-clustered standard errors.

### Software

Analyses used Python with pandas, NumPy, scikit-learn, XGBoost, statsmodels, SHAP, matplotlib and seaborn. The pipeline is organised as a modular project with a single parameter file, and every entry point takes the input, output and split-year settings from the command line so that each organism runs without code changes.

## Supporting information

Supplementary_Data.zip: Supplemental Data (replicate-level results)

