## Supplementary_Data.zip: Supplemental Data (replicate-level results) for "Forecasting next-year change in antimicrobial resistance genes in *Klebsiella pneumoniae* and *Acinetobacter baumannii* from genomic surveillance data": README.txt

Replicate-level results for the gene-year forecasting study (K. pneumoniae = KP, A. baumannii = AB).

*_replicate_level_results.csv
  One row per model configuration and replicate (seed 0 to 24). Columns: target (emergence or turnover), seed,
  split (temporal or gene_held_out), model, feats (stability index variant; "-" for baselines), AUROC, AUPRC,
  threshold-based metrics (precision, recall, f1, specificity, npv, accuracy, bal_accuracy, mcc), brier,
  threshold (classification threshold selected on training data), prevalence (event rate in the test data),
  and confusion-matrix counts (tp, fp, tn, fn).

*_replicate_level_SI_ablation.csv
  One row per model, outcome, evaluation scheme, stability index variant and replicate. Columns: AUROC and AUPRC
  with and without the stability index, and the paired differences d_auroc and d_auprc.

Summaries of these files are in Supplementary_Tables.xlsx.
