## Supplementary_Figures.pdf: Supplemental Figures S1 and S2 for "Forecasting next-year change in antimicrobial resistance genes in *Klebsiella pneumoniae* and *Acinetobacter baumannii* from genomic surveillance data"

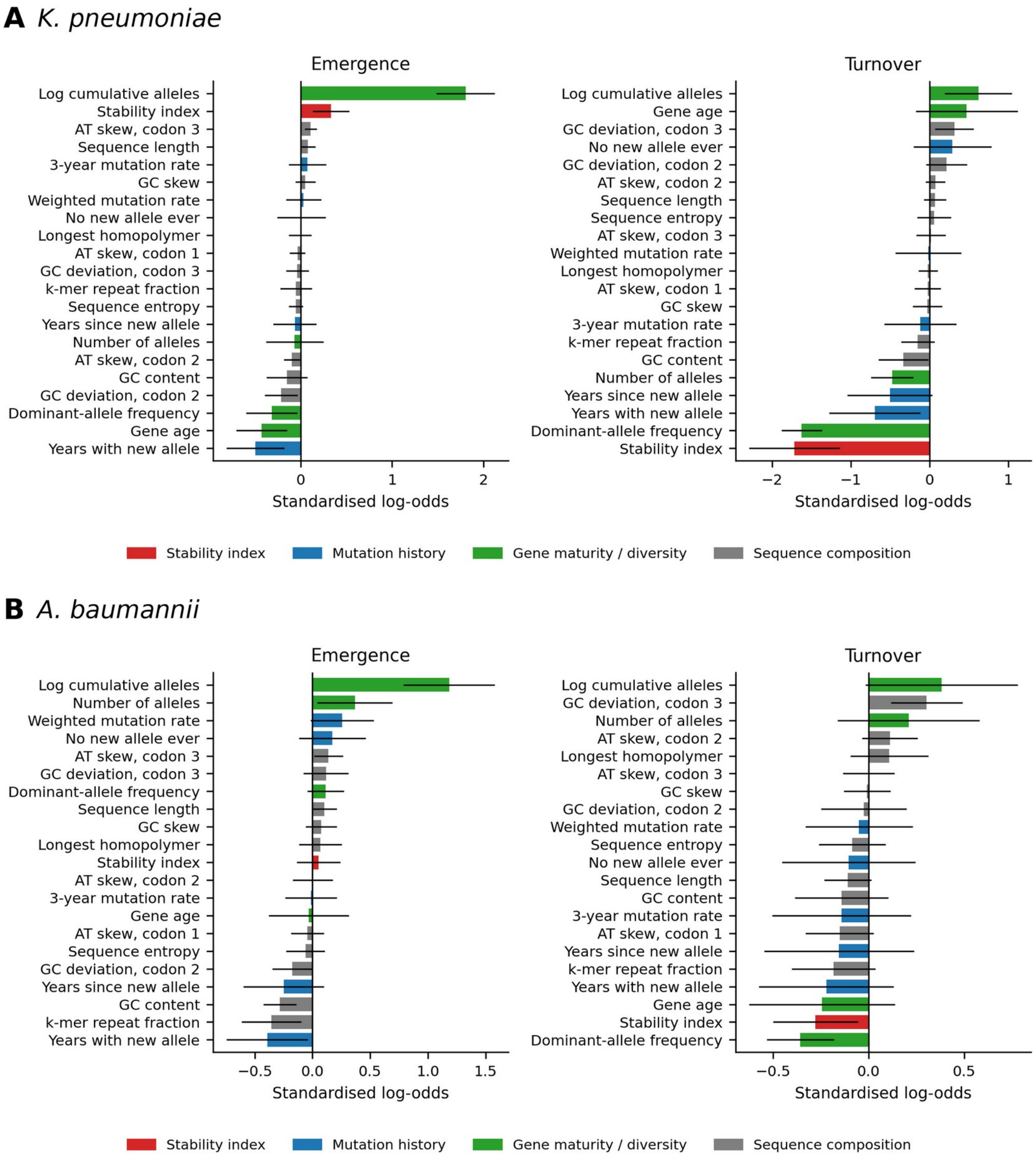


**Figure S1. Logistic regression coefficients.** Mean standardised log-odds coefficient for each feature, for emergence (left) and turnover (right). A, *K. pneumoniae*; B, *A. baumannii*. Bars are coloured by feature block, and error bars span 1.96 standard deviations of the coefficient across 25 replicates (not a confidence interval). The three missingness indicators are not shown because their estimates are unstable when the indicator is rare (see Table S4 for all odds ratios).

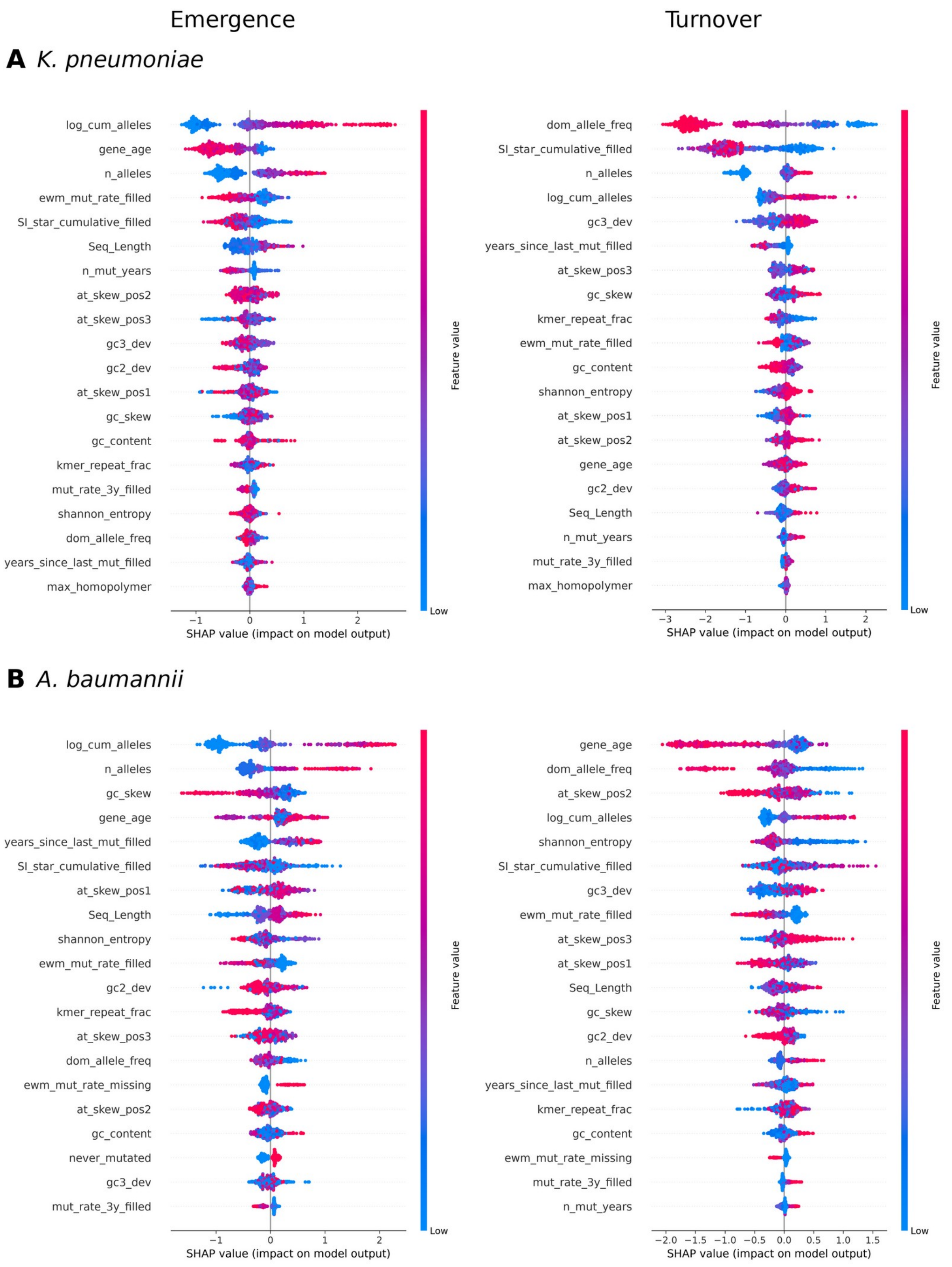


**Figure S2. SHAP values for individual gene-years.** Each point is a gene-year in the test years. Its position shows the SHAP value (impact on the model's log-odds output) and its colour the value of the feature (blue low, red high). The 20 features with the largest mean absolute SHAP value are shown, from an XGBoost model fitted to the training years of replicate 0, for emergence (left) and turnover (right). A, *K. pneumoniae*; B, *A. baumannii*.
